# Correlations of genotype with climate parameters suggest *Caenorhabditis elegans* niche preferences

**DOI:** 10.1101/075960

**Authors:** Kathryn S. Evans, Yuehui Zhao, Shannon C. Brady, Lijiang Long, Patrick T. McGrath, Erik C. Andersen

## Abstract

Species inhabit a variety of environmental niches, and the adaptation to a particular niche is often controlled by genetic factors, including gene-by-environment interactions. The genes that vary in order to regulate the ability to colonize a niche are often difficult to identify, especially in the context of complex ecological systems and in experimentally uncontrolled natural environments. Quantitative genetic approaches provide an opportunity to investigate correlations between genetic factors and environmental parameters that might define a niche. Previously, we have shown how a collection of 208 whole-genome sequenced wild *Caenorhabditis elegans* can facilitate association mapping approaches. To correlate climate parameters with the variation found in this collection of wild strains, we used geographic data to exhaustively curate daily weather measurements in short-term (three month), middle-term (one year), and long-term (three year) durations surrounding the data of strain isolation. These climate parameters were then used as quantitative traits in the mapping approaches. We identified 10 QTL underlying variation in three traits: elevation, relative humidity, and average temperature. We then performed statistical analyses to further narrow the genomic interval of interest to identify gene candidates with variants potentially underlying phenotypic differences. Additionally, we performed two-strain competition assays at high and low temperatures to validate a QTL for temperature preference and found suggestive evidence that genotypes might be adapted to particular temperatures.

**100-word summary for *G3*:** Quantitative genetic approaches provide an opportunity to investigate correlations between genetic factors and environmental parameters that might define a niche, but these genes are difficult to identify, especially in the context of complex ecological systems. Here, we used a collection of 152 sequenced wild *Caenorhabditis elegans* to correlate climate parameters with the variation found in this collection of wild strains. We identified 10 QTL in five traits, including elevation, relative humidity, and temperature. Additionally, we performed competition assays to validate a QTL for temperature preference and found suggestive evidence that genotypes might be adapted to particular temperatures.

## INTRODUCTION

Ecological niches describe how individuals of a species respond to and alter the distribution of resources and competitors within their environment (Hutchinson 1957). These resources could include food availability, soil type, short-term weather conditions, and long-term climate. Often, a species can be found in multiple distinct geographic areas that all share a common set of environmental factors and resources. For example, a plant that thrives at high temperature might grow equally well anywhere along the equator. An organism’s ability to survive in a specific niche is driven by both environmental and genetic factors. Genetic variation between species, and among individuals within a species, contributes to the wide variety of niches observed (Saltz and Nuzhdin 2014). A genetic variant could result in an increased affinity for an individual to its environment. This individual will be selected and, over time, evolution will favor the successful variant. This phenomenon, known as gene-by-environment interactions, refers to when different genotypes respond to environmental variation in diverse ways.

Previous studies in model organisms, particularly *Drosophila*, have investigated gene-by-environment interactions with clinal variation. Selection on body size is correlated with temperature (Taylor *et al.* 2015), and survival is affected by climate change (Bozinovic *et al.* 2016). Machado *et al.* performed a longitudinal study of *Drosophila* collected at differing latitudes during a two-year time span and compared physiological traits of two different species – *D. melanogaster* and *D. simulans* (Machado *et al.* 2016). Other studies have gone further by identifying quantitative trait loci (QTL) for body size and cold tolerance traits involved in adaptation to seasonally varying environments (Tyukmaeva *et al.* 2015; Hangartner *et al.* 2015). Gerken *et al.* found substantial heritable variation in both short-term and long-term acclimation (Gerken *et al.* 2015). They then performed genome-wide association mappings on these traits and found the QTL for short-term and long-term adaptations did not overlap, but each resulted in a set of gene candidates sharing similar functions of apoptosis, autophagy, cytoskeletal and membrane structural components, and ion binding and transport. Missing from these studies is genome-wide association mappings for other environmental traits besides temperature and latitude. Furthermore, little has been studied for other model organisms, such as *Caenorhabditis elegans*.

*C. elegans* is a free-living nematode often found in microorganism-rich organic material such as rotting fruits and compost heaps in temperate and humid environments (Frézal and Félix 2015; Félix and Braendle 2010). The first studied *C. elegans* strain, N2, was isolated from mushroom farm compost in Bristol, England in 1951 (Hodgkin and Doniach 1997). Since that time, N2 has been used as the wild-type strain for *C. elegans* laboratory research. This strain was cultured for many years in the laboratory, potentially resulting in selection for alleles favorable in that environment (Sterken *et al.* 2015). To study natural variation and the role of niche specification on this species, we require a worldwide collection of wild strains. Our research group acquired a large collection of 208 wild strains and sequenced the whole genomes of these strains (Cook *et al.* 2016). By comparing the genomes of the 208 strains, we found that some strains from similar geographic locations have nearly identical genome sequences. This analysis results in 152 unique genome-wide haplotypes or isotypes. This large pool of genetic information provides us with the statistical power to make connections between genotype and phenotype using genome-wide association (GWA) studies. GWA studies use common genetic variation across a population of individuals to discover variants that are correlated with a specific trait of interest (Hirschhorn and Daly 2005).

In this study, we correlate natural genetic variation among 152 wild *C. elegans* strains with climate measurements of their environmental niches as quantitative traits. We mapped traits that describe the niche of the isolation location for each strain, including geographic parameters, seasonal weather patterns, and climate variables. We find significant associations for elevation, relative humidity, and temperature. These findings suggest genetic control of niche specification. Additionally, we tested the quantitative trait locus (QTL) associated with temperature and found possible evidence of a specific preference for lower temperatures based on genetic background.

## RESULTS

### Genome-wide association of geographic traits

The location where a *C. elegans* strain was identified could reflect the process of selection for a particular genotype in a specific niche. For this reason, we investigated correlations between genetic variation in the *C. elegans* population and parameters describing the geographic locations of isolation as quantitative traits. Previous work with a smaller set of strains (97 wild isolates) detected a significant QTL on the left arm of chromosome II associated with the latitude where strains were isolated (Andersen *et al.* 2012). To evaluate this trait and other parameters describing the location where each strain was isolated in our larger strain set (152 strains), we curated the isolation location information and defined several traits based on geographic data for each strain (Table 1), namely latitude, longitude, elevation, the absolute value of latitude, and the absolute value of longitude (File S1). We performed genome-wide association mappings with 149 wild strains with known isolation locations (Figure 1A) to correlate these trait values with common genetic variation (File S2; *see Methods*). Using this strain set, only the mapping of the elevation of the isolation location identified a significant QTL on the left arm of chromosome III (Figure 2A). When we divided the population by the genotype at the peak marker, we found that the elevation values for these two sets of strains were similar with a few outliers (Figure 2B), suggesting that the outliers were causing the detection of a QTL. It is possible that the association is spurious and driven by outliers or that the outliers are strains harboring rare alleles in the *C. elegans* species that impact this trait. The outlier strains in our mapping could share some genetic similarities as they were all collected within the last 15 years, and most were collected in France or elsewhere in Northern Europe. We did not recapitulate the QTL for latitude observed in the previous study (Andersen *et al.* 2012) likely because it also appears to be driven by strains with extreme latitude values. Again, these outlier strains could be highly related as seven of the 15 strains with the alternate genotype at the peak marker position originated from South Africa or Kenya. Our larger strain set reduces the effect of these outliers on the GWA mapping, and the previously detected QTL is no longer significant. These results suggest that common variation in the *C. elegans* species does not correlate with geographic parameters describing the location of strain isolation. However, rare variants might control whether strains can colonize and/or proliferate in specific geographic locations.

**Figure 1.**
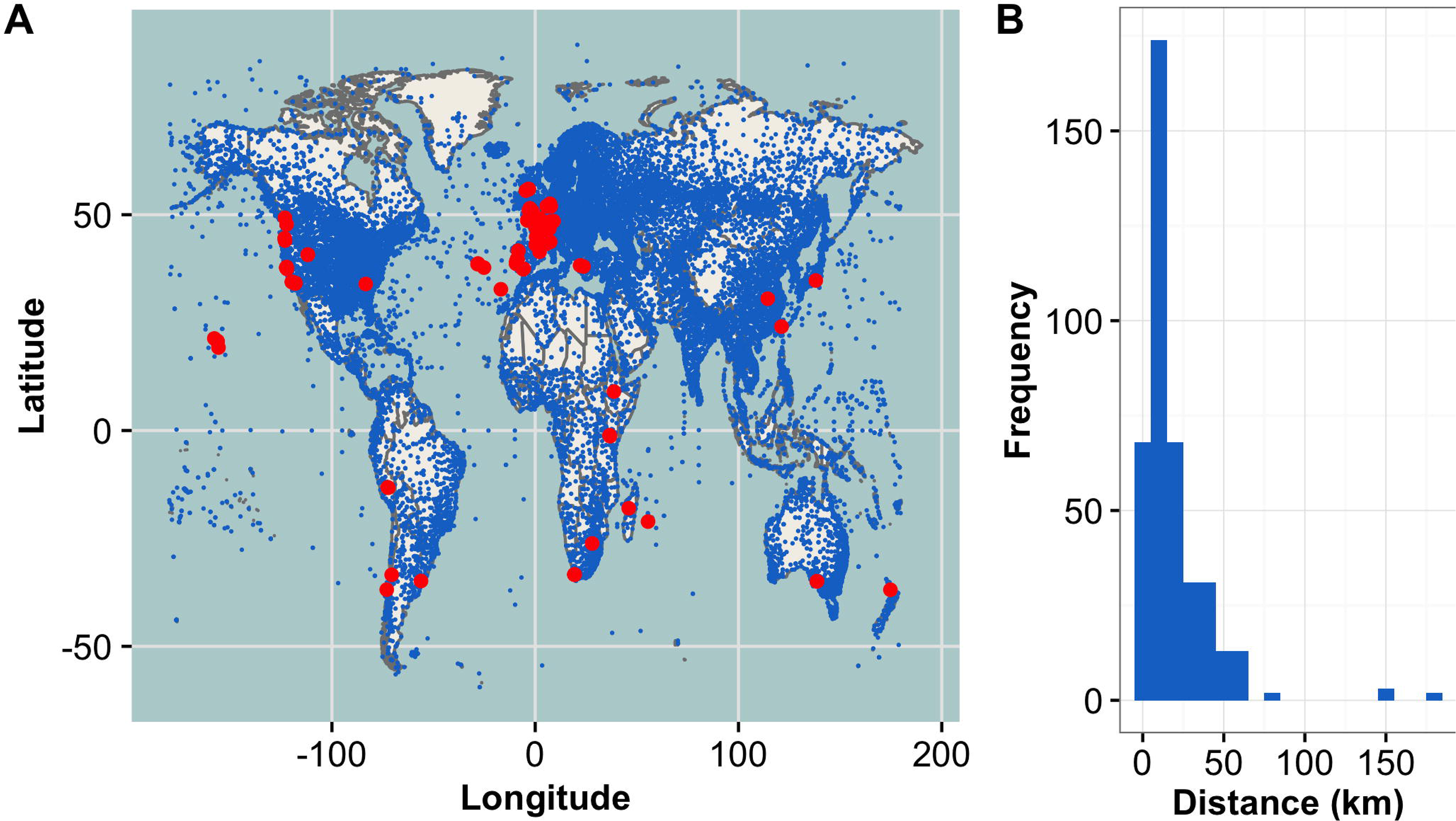
Global distribution of wild isolates and NOAA weather stations. A) Map of 27,447 ISD NOAA weather stations (blue) and 149 *C. elegans* wild isolate locations (red). Three isotypes are not depicted as they have no known location of isolation. B) Histogram of station distance from wild isolate, measured in kilometers (km). Data from the three-year weather dataset are shown.

**Figure 2.**
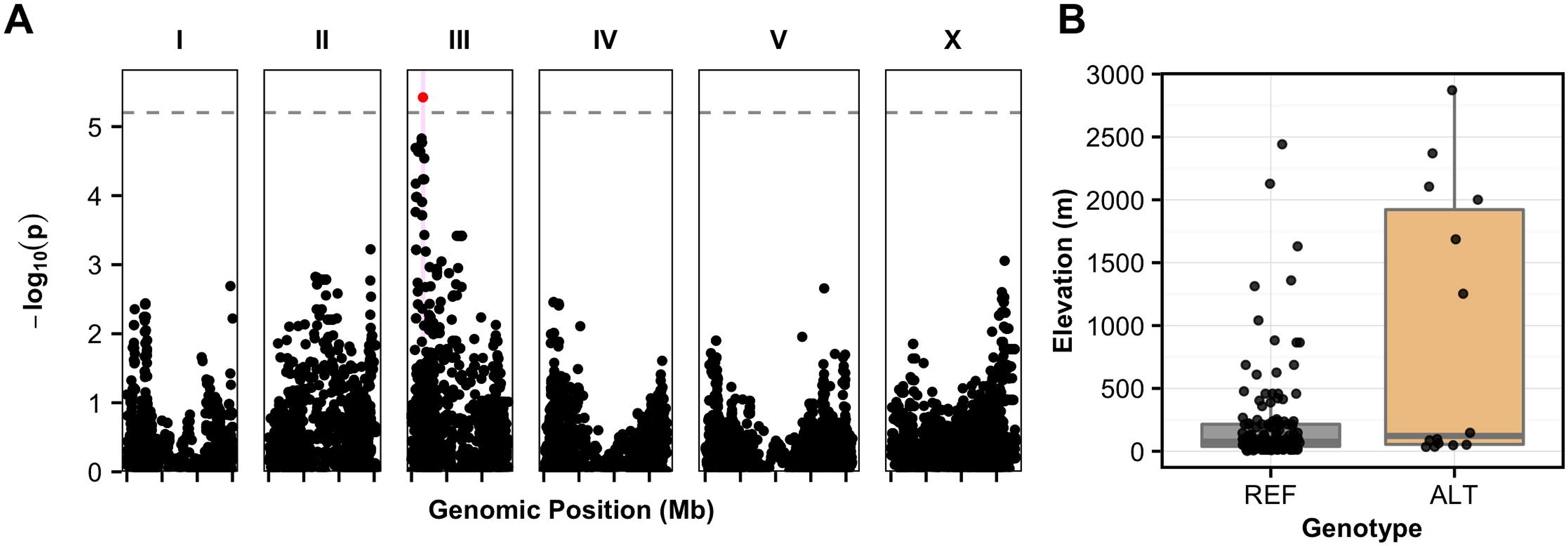
Genome-wide association of elevation. A) Genome-wide association of elevation of strain isolation shown as Manhattan plots. Genomic position is plotted on the x-axis against the negative log-transformed *p*-value on the y-axis. Single nucleotide variants (SNVs) that are above the Bonferroni-corrected significance threshold, indicated by the dotted grey line, are shown in red and SNVs below the Bonferroni threshold are shown in black. Confidence intervals are represented by the pink bars. B) Box plots show strain isolation elevations, measured in meters (m), separated by the genotype at the peak marker location. Each point represents one strain. The reference genotype (REF) refers to strains that share the genotype of the reference strain, N2. The alternative genotype (ALT) refers to all other strains that do not have the reference genotype at the peak marker position.

### Weather conditions and climate parameters can be determined using the geographic location of the site of strain isolation

It is likely that the possible genetic association we observed between the elevation of strain isolation and a region on chromosome III is correlated with weather patterns and/or climate variables at specific geographic locations. Strain latitude and longitude can be used to determine the weather or climate at the time and location from which each wild *C. elegans* strain was isolated. The short-term weather surrounding the day of isolation as well as the long-term climate of the geographic location for each strain could improve our understanding of possible preferred niches for specific strains or for the species as a whole. Furthermore, combining these climate parameters with whole-genome sequence data could identify potential alleles that underlie preferences for certain environmental factors.

The National Oceanic and Atmospheric Administration (NOAA) collects and provides multiple datasets related to weather and climate information, including the Integrated Surfaces Data (ISD). The ISD dataset is archived at the National Climatic Data Center (NCDC) and is composed of worldwide surface weather observations from over 27,000 stations managed by different global institutions (“Integrated Surface Global Hourly Data - NOAA Data Catalog” 2016). First, we manually curated the isolation information for the 152 strains, including the date of isolation, location, and sampling information (File S1). Then, we overlaid the locations of the 27,447 ISD weather stations with the isolation locations of the 149 *C. elegans* wild strains with complete sampling data (Figure 1A). Using the date and isolation location for each wild strain, we identified the closest weather station with available data and collected weather observations in three time periods surrounding the date of nematode isolation: three months, one year, and three years. Most strains were found less than 60 km from a weather station (Figure 1B). Of the 149 strains with known isolation locations, we knew at least the year of isolation for 145 strains, the month for 138 strains, and the day for 122 strains. For the three-month period, we analyzed weather station data only for strains with known days of isolation to provide a precise account of the daily weather experienced immediately surrounding the date of isolation for each strain. However, for the one-year period we used data from strains with known day or month of isolation. For the three-year period, we used data from strains where either the day, month, or year of isolation is known to provide an estimated overall climate of the strain isolation location. Not every weather station sampled contained data for each weather parameter. Additionally, only quantitative weather parameters that were measured at locations shared in a majority of the wild-isolate population (more than 90% of the strains) were considered for further analysis (Table 1). All observations for each weather parameter were averaged over the given time period, and this average value was used as the trait measurement for each strain (File S2, *see Methods*).

**Table 1.** Definition of geographic and weather traits ^a^

| Trait name | Trait code<sup>b</sup> | Number of strains<sup>c</sup> | Description |
| --- | --- | --- | --- |
| **Latitude** | latitude | 149 | Latitude coordinate where the nematode strain was collected (degrees) |
| **Abs. value of latitude** | abslat | 149 | Absolute value of the latitude coordinate where the nematode strain was collected (degrees) |
| **Longitude** | longitude | 149 | Longitude coordinate where the nematode strain was collected (degrees) |
| **Absolute value of longitude** | abslong | 149 | Absolute value of the longitude coordinate where the nematode strain was collected (degrees) |
| **Elevation** | elevation | 149 | Calculated elevation based on latitude/longitude coordinates using “geosphere” package (m) |
| **Temperature** | temp | 145 | Temperature of the air (°C) |
| **Relative humidity** | rh | 129 | Amount of water vapor present in air expressed as a percentage of the amount needed for saturation at the same temperature (%) |
| **Wind direction** | wd | 134 | Angle, measured in a clockwise direction, between true north and the direction from which the wind is blowing (angular degrees) |
| **Wind speed** | ws | 136 | Rate of horizontal travel of air past a fixed point (m/s) |
| **Cloud height<sup>d</sup>** | ceil_hgt | 122 | Height above the ground level of the lowest cloud or obscuring phenomena layer with 5/8 or more summation total sky cover, which may be predominantly opaque, or the vertical visibility into a surface-based obstruction (m) |
| **Dew point** | dew_point | 129 | Temperature to which a given parcel of air must be cooled at constant pressure and water vapor content in order for saturation to occur (°C) |

We evaluated all weather observations over the three months, one year, or three years surrounding the date of nematode isolation for each of the 149 wild strains to define the weather or climate experienced by each strain. These data were mapped using GWA to define nine QTL (Figure S1) for two distinct traits – relative humidity and temperature.

### Genome-wide association of average relative humidity

*C. elegans* is found at various relative humidities ranging from 36% to 92%, with an average of 71.8% (File S2). This estimate of the average relative humidity of the *C. elegans* niche is in concordance with previous studies that show *C. elegans* is often found in humid environments (Frézal and Félix 2015). To determine if variation in relative humidity at isolation location is correlated with genetic variation, we performed GWA mapping for relative humidity of isolation location for three months, one year, and three years surrounding the date of isolation (*see Methods*). We found eight significant QTL in three distinct regions of the *C. elegans* genome: the left arm of chromosome II, the right arm of chromosome III, and the right arm of chromosome V (Figure 3, File S2). The mapping for average relative humidity over three months surrounding the date of isolation (Figure 3A) resulted in two linked QTL on the left arm of chromosome II (LD r^2^ = 0.999, Figure S1) and a QTL on the right arm of chromosome III that is also in high linkage disequilibrium with the QTL on chromosome II (LD r^2^ = 0.634, 0.812; Figure S1). The mapping for average relative humidity over one year surrounding the date of isolation (Figure 3B) resulted in one QTL on the left arm of chromosome II and two highly linked QTL on the right of chromosome V (LD r^2^ = 0.536; Figure S1). The position of the chromosome II QTL is the same as that observed for the mapping of three-month humidity. Finally, the mapping of average relative humidity over the three years surrounding the date of isolation (Figure 3C) resulted in the same two QTL on chromosome V as found for the mapping of one-year relative humidity (Figure S1). For each QTL, strains with the reference allele at the peak marker tend to be isolated at higher relative humidities, and strains with an alternative allele tend to be isolated at lower relative humidities (Figure S1). This evidence of a phenotypic split dependent on genotype of the peak marker suggests that at least one variant could underlie a preference for relative humidity.

**Figure 3.**
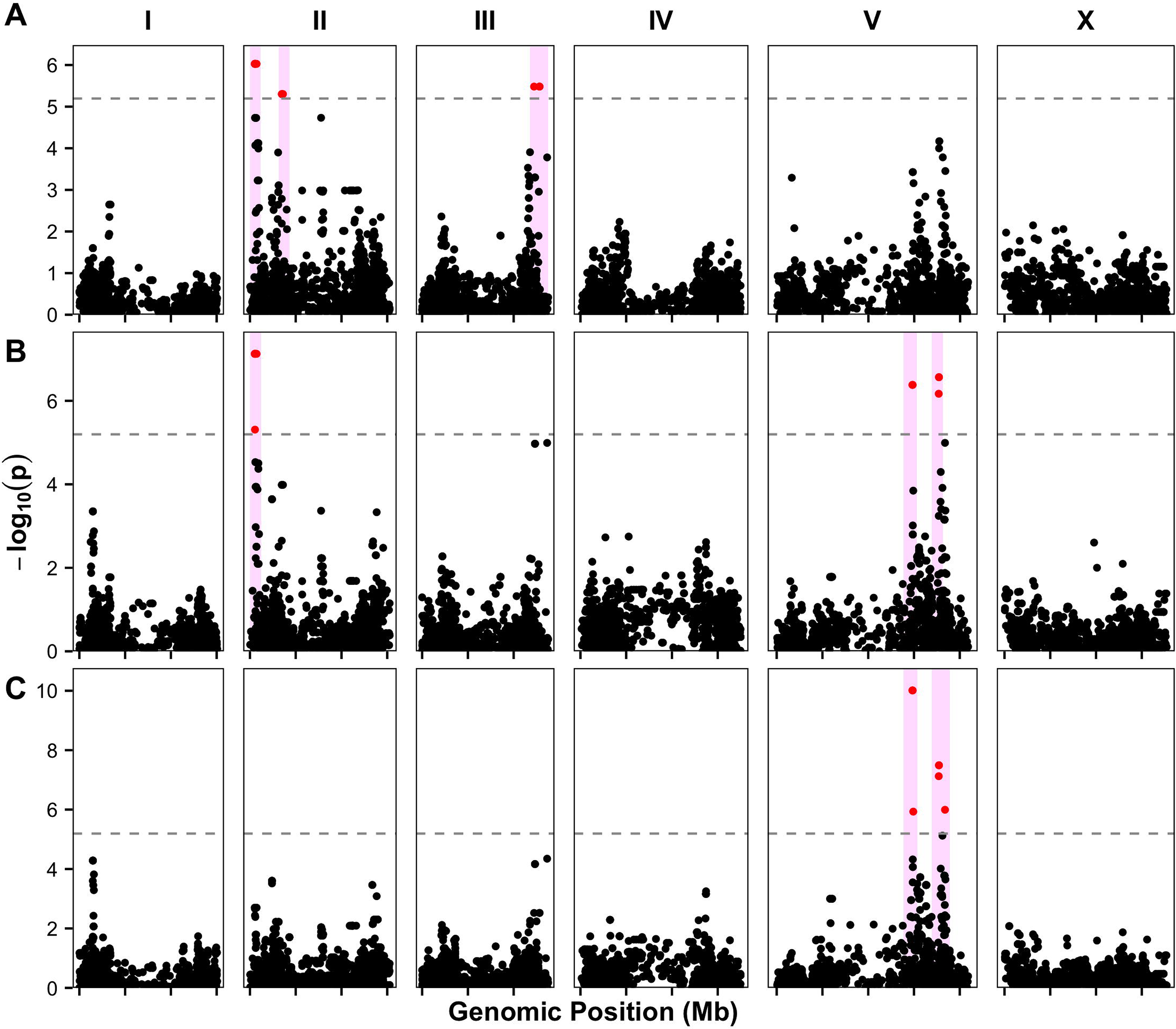
Genome-wide association of humidity traits. Genome-wide association of relative humidity for three different time periods are visualized as Manhattan plots: three-month (A), one-year (B), three-year (c) durations. Genomic position is plotted on the x-axis against the negative log-transformed *p*-value on the y-axis. SNVs that are above the Bonferroni-corrected significance threshold, indicated by the dotted grey line, are shown in red and SNVs below the Bonferroni threshold are shown in black. Confidence intervals are represented by the pink bars.

### Genome-wide association of average temperature

*C. elegans* are also found at a variety of average temperatures ranging from 7°C to 25°C, with an average of 15.3°C (File S1). To determine if temperature is associated with genetic variation, we performed a GWA mapping for temperature of isolation location for three months, one year, and three years surrounding the date of isolation (*see Methods*). We found one significant QTL just right of the center of chromosome V (Figure 4A; File S2). This QTL is in the same location as that observed for one-year and three-year relative humidity (Figure 3B, 3C). We found that strains with the reference (N2) allele at the peak marker tend to be isolated from geographic locations with lower temperatures, and strains with an alternative allele at this position tend to be isolated from geographic locations with higher temperatures (Figure 4B). This QTL suggests that an allele, or alleles nearby this marker, could confer survival advantages to strains that experience different temperatures. To identify the variant(s) that could be driving this QTL, we investigated a region on chromosome V (V:13,845,281-15,332,878) defined by 1.48 Mb that contains 619 total genes. The genes with predicted functional variants are most likely to cause phenotypic differences among diverse strains in species. Therefore, we focused on 363 genes within this region predicted to harbor functional variants of moderate or severe effects on gene function, as determined by SnpEff (Cingolani *et al.* 2012). We can further narrow our list of candidate genes to 48 by identifying genes that are highly correlated with differences in temperature (File S3). Although an investigation of these 48 genes did not identify an obvious candidate related to temperature regulation, one or more of these genes could be responsible for the preference of certain strains for specific temperatures.

**Figure 4.**
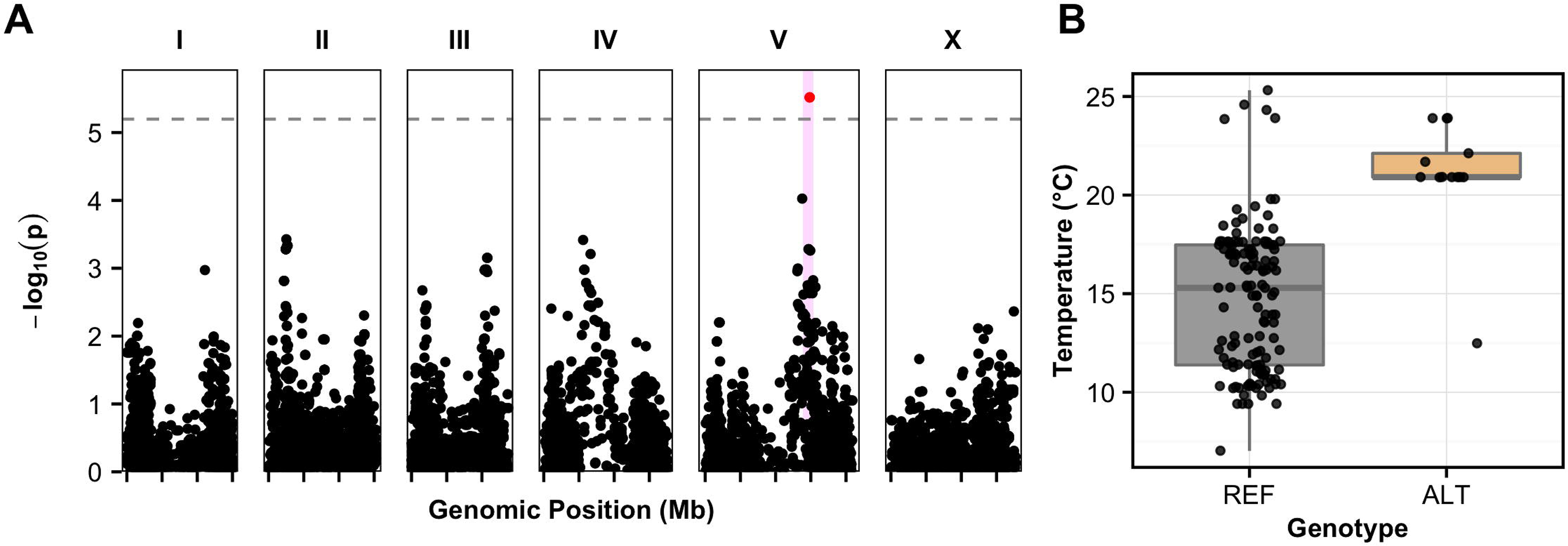
Genome-wide association of temperature. A) Genome-wide association of three-year average temperature is visualized as a Manhattan plot. Genomic position is plotted on the x-axis against the negative log-transformed *p*-value on the y-axis. SNVs that are above the Bonferroni-corrected significance threshold, indicated by the dotted grey line, are shown in red, and SNVs below the Bonferroni threshold are shown in black. Confidence intervals are represented by the pink bars. B) Box plots show the strain three-year average temperatures, measured in degrees Celsius, separated by genotype at the peak marker locus. Each point represents one strain. The reference genotype (REF) refers to strains that share the genotype of the reference strain, N2. The alternative genotype (ALT) refers to all other strains that do not have the reference genotype at the peak marker position.

### Strains from divergent climates might be adapted to specific temperatures

Although we have GWA mappings for various weather conditions, validating these QTL would provide more evidence for *C. elegans* selection of niche based on environmental and geographic factors. Because temperature can be controlled easily and survival at defined temperatures can be tested experimentally, we decided to determine whether two strains from divergent climates are adapted to the respective temperatures nearby their isolation locations using a competition assay. Strains were chosen that had both a different genotype at the peak marker of the QTL on chromosome V identified in the three-year temperature mapping experiment and a large difference in the temperatures nearby the isolation location. JU847, a strain isolated from Northern France in 2005 (File S1), has the reference genotype at the three-year temperature QTL peak marker and is found at a low three-year average temperature (11.3°C). CX11314, a strain isolated from Southern California, USA in 2003 (File S1), has the alternative genotype at the three-year temperature QTL peak marker and is found at a higher three-year average temperature (20.9°C). High fitness at or around the temperature nearby the isolation location of one strain and lower fitness at or around the temperature nearby the isolation location for the other strain would suggest potential adaptive alleles that contribute to a preference and/or better survival at a specific temperature.

Replicate cultures were initiated with an equal number of animals from each strain at either 15°C or 25°C and allowed to compete for at least six generations. After culture transfers two, four, and six, we analyzed the ratios of the two strains found at each temperature (*see Methods*). CX11314 was found to have higher fitness than JU847 at both temperatures tested (Figure 5). At high temperature, CX11314 had a clear selective advantage compared to JU847 (for fitness = 1, relative CX11314 fitness s = 2.29), resulting in JU847 alleles comprising less than 1% of the alleles measured after six culture transfers. However, JU847 performed better at the lower temperature than at the higher temperature, comprising almost 8% of the total nematode population after six culture transfers (relative CX11314 fitness s = 1.57). These data suggest that JU847, although not more fit than CX11314 at either temperature, is more fit at 15°C than at 25°C.

**Figure 5.**
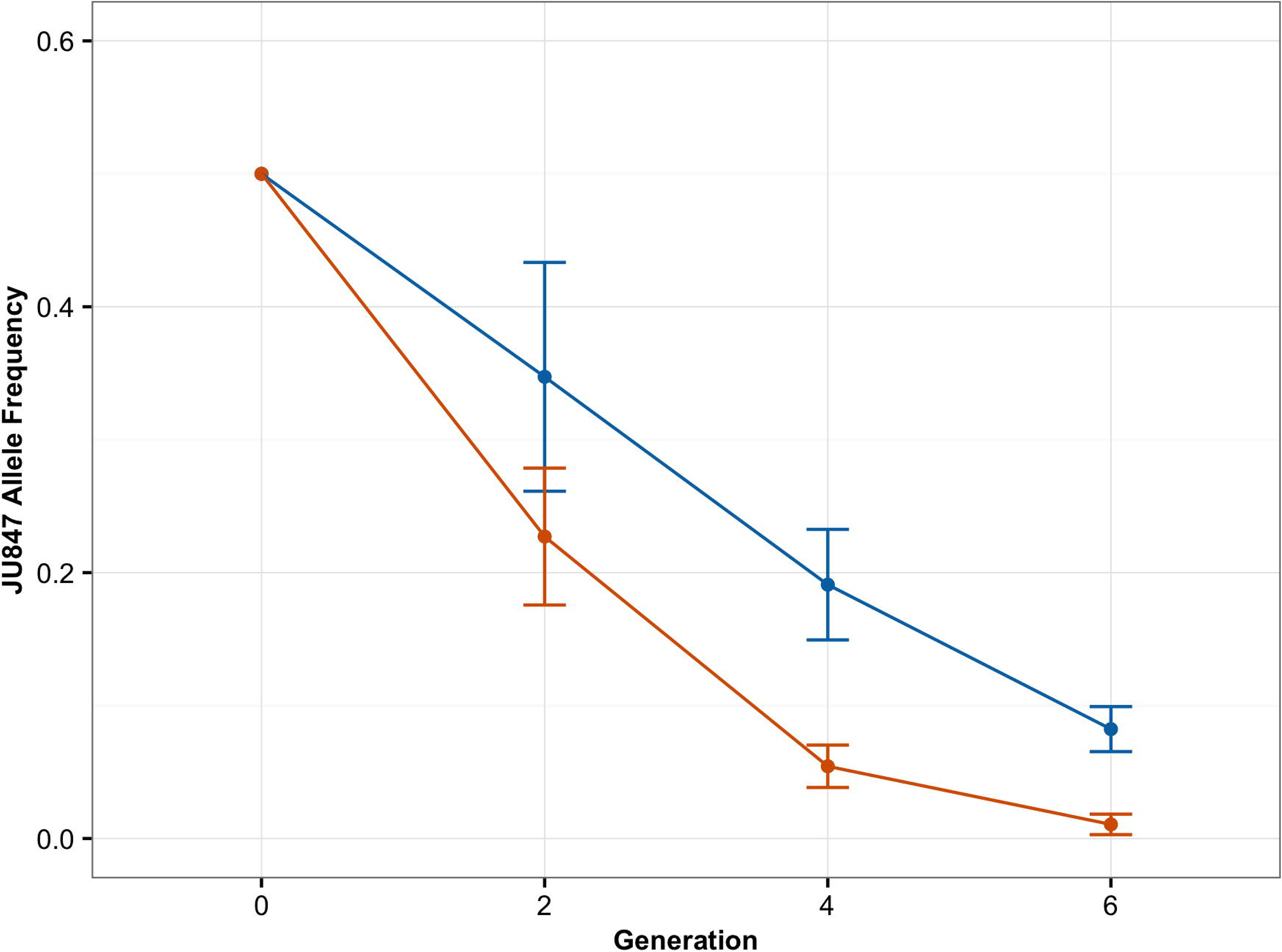
Two-strain temperature competition assay. JU847 (isolated at low temperature) was competed against CX11314 (isolated at high temperature) at both 15°C (indicated in blue) and 25°C (indicated in red). The mean frequency of the JU847 allele in the population is plotted on the y-axis. Error bars represent one standard deviation from the mean. Data were collected from nine experimental and five technical replicates.

## DISCUSSION

In this study, we have defined geographic, weather, and climate variables over three different time periods as phenotypic traits for 149 unique wild *C. elegans* strains and performed genome-wide association mapping for 11 traits. Each phenotype described in this study was obtained using the location, date, and weather of the known isolation location of each isotype in our collection. We found significant correlations between genotype and phenotype for three traits and a total of 10 QTL. However, only temperature displayed strong phenotypic separation associated with genotypic variation that was likely not driven by outlier strains.

Although we found a significant QTL associated with the elevation of isolation location, we did not recapitulate the QTL for latitude observed in the previous study (Andersen *et al.* 2012). This difference is likely because the previous mapping appears to be driven by strains with extreme latitude values, similar to our elevation mapping. The larger strain set in this analysis reduces the effect of these outliers on the GWA mapping, and thus the QTL is no longer significant. However, we did observe a QTL associated with latitude just below the Bonferroni-corrected significance threshold on the right arm of chromosome V. This suggestive QTL is in close proximity to the QTL for temperature. It is possible that the moderate association seen with latitude is representative of a real association with temperature, as the two traits are highly correlated. In our dataset, we see a high negative correlation between the absolute value of latitude and temperature (*rho* = −0.805), and between latitude and temperature (*rho* = −0.772). This lower-than-expected correlation is likely due to temperature fluctuations throughout the year: temperature at high latitude in the summer would be similar to temperature near the equator in the winter. Additionally, the latitude and longitude recorded for a particular strain is not always precise, especially for strains with older isolation dates. One tenth of a degree of latitude can distinguish between large cities, but could cover up to 11.1 km of distance. The function we used to determine elevation of strain isolation used these imprecise latitude and longitude coordinates, potentially resulting in a range of small to sizeable errors. These estimations could affect not just the geographic traits, but the weather traits as well, which are calculated based on the geographic coordinates. However, we do not expect this estimation to have a large effect, as we found weather data between stations up to 150 km apart to be highly correlated (*see Methods*).

We chose to assess weather conditions for each of the 149 strains over three time periods: three months, one year, and three years. A duration of three months was chosen to define the weather contemporary to the date of isolation. One year was chosen to evaluate the average weather patterns a strain must be able to survive in nature. Three years was chosen to define the climate of the isolation location. Using data from three years could help us understand the average long-term climate conditions for each strain by eliminating any unusual weather patterns during the year of isolation that are not representative of the overall average environmental conditions. The relative humidity trait maps to the same chromosome positions for all three time periods assayed, suggesting that relative humidity remains constant throughout seasons. Previous studies have also shown that *C. elegans* tend to be found in humid regions (Frézal and Félix 2015). The same QTL on the right arm of chromosome V was observed for both long-term relative humidity and long-term temperature. Because relative humidity depends on air temperature, these results are expected to be correlated. Although the average temperature maps with only the three-year data set, we observed a QTL at the same position just below the significance threshold for the one-year mapping dataset as well. This suggestive QTL provides more evidence that one or more variants on chromosome V are associated with differences in long-term average temperature of isolation for *C. elegans*.

The QTL for temperature was evaluated further because it could be controlled in a laboratory setting. The competition assay between JU847 and CX11314 showed that CX11314 had a higher selective advantage than JU847 at both high and low temperatures. This result could be observed because the laboratory environment can not completely recapitulate conditions experienced in the wild. Additionally, it is possible that CX11314 is more fit than JU847 regardless of temperature. To test this QTL more thoroughly and eliminate this possibility, it would be necessary to compete multiple high and low temperature strains. We found that the low temperature strain, JU847, was more competitive with the high temperature strain, CX11314, at the lower temperature. This result provides evidence that one or more genetic variants within the QTL could underlie the hypothesized temperature preference. Furthermore, the duration of this experiment was only six weeks. At the lower temperature, it is possible we could observe a stronger competitive advantage for JU847 over a longer time period.

Although our analyses were unable to identify a single gene or variant that could underlie potential differences in niche specification, our conclusions suggest that different strains are found in unique niches and at least some of the environmental differences in niches are related to genetic variation among strains. As we expand our collection of wild *C. elegans* strains, we will be able to better define these weather and climate differences. Additionally, longitudinal collection studies with dense sampling, especially in a location with known high species diversity (such as the Hawaiian Islands) would give us more valuable data about how genetic variation in *C. elegans* is related to environmental conditions. We expect that similar data could be analyzed for other species and allow for investigation of niche specification, specifically in plant species where dense sampling and whole-genome datasets are available.

## MATERIALS AND METHODS

### *C. elegans* wild isolate collection and sequencing

A collection of 208 wild *C. elegans* strains have been isolated worldwide and annotated for each strain’s geographic location and date of isolation. Members of the Andersen Lab have carefully and manually curated this dataset to offer the most accurate information possible while accounting for sometimes imprecise sample recording. Whole-genome sequence (WGS) data were collected from all 208 strains (Cook *et al.* 2016). The raw Illumina data are deposited with the Short Read Archive under project PRJNA318647. WGS data were analyzed as previously described. In brief, after alignment with BWA (Li and Durbin 2009) and variant calling using Samtools (Li *et al.* 2009), strains with a concordance of 99.93% or higher were grouped as a genome-wide haplotype or isotype. This analysis resulted in 152 unique isotypes (File S1).

### Weather and climate data acquisition

For each wild strain with a known isolation location, elevation was estimated with the “geosphere” package in R (Hijmans, Williams, and Vennes 2012) using the geographic coordinates of strain isolation. A correlation test using 74 points of known elevation were used to verify accuracy of the elevation function resulting in a correlation of 0.998. Weather data were downloaded from the Integrated Surface Data (ISD) FTP server (ftp://ftp.ncdc.noaa.gov/pub/data/noaa/) managed by the National Oceanic and Atmospheric Administration (NOAA) and the National Climatic Data Center (NCDC). The ISD dataset comprises worldwide surface weather observations from over 27,447 stations managed by the Automated Weather Network (AWN), the Global Telecommunications System (GTS), the Automated Surface Observing System (ASOS), and others. Data are collected once every three hours for some stations. Some parameters include air quality, atmospheric pressure, atmospheric temperature, dew point, atmospheric winds, clouds, precipitation, ocean waves, and tides.

Three distinct sets of weather station data were collected for analysis: a three-month window, a one-year window, and a three-year window. These data were filtered to include values centered around the date of isolation. For the three-month set, wild isolates with only a known month or year of isolation were not considered. Exact day of isolation is necessary to understand the seasonal environment in which an animal was isolated. For the one-year dataset, wild isolates with a month or day of isolation were used. For the three-year dataset, strains with only a year of isolation were used in addition to those strains with more defined dates of isolation. If only the year was known, the date of isolation defaulted to January 1 of that year, and data were collected surrounding that date. If only the month of isolation was known, the date of isolation was defaulted to the first of that month for data collection.

The 27,447 NOAA weather stations were filtered by their availability of data collected within the years of interest. Stations that had less than ten recordings of any type for any month within the time period of data collection were excluded to avoid misrepresentation by datasets that were averaged from only a few data points. Stations were then filtered by location, and the closest station to location of isolation for each wild strain was selected and downloaded using the “stationaRy” package available at https://github.com/rich-iannone/stationaRy (Iannone 2015). We performed a rank-correlation test for temperature, relative humidity, and atmospheric pressure between two neighboring weather stations (ranging from 0.93-153 km apart) and found high correlations regardless of distance between stations (*rho* = 0.920, 0.913, and 0.707, respectively). All station-isotype pairs were included in our analysis regardless of the distance between them. The primary fields as well as all additional quantitative data available for each station were downloaded. Some fields (*e.g.* “AT1” or “Present-weather-observation”) were not downloaded from the NOAA database because the traits are qualitative and would not be conducive to quantitative analyses. The station data were filtered to contain only information from the months surrounding the date of isolation. This process was repeated for each dataset (three-month, one-year, and three-year) in case a closer weather station contained only data for the three-month set but not for the one-year or three-year sets. The station data were meticulously checked by manually removing missing values from each weather category independently and, in certain cases, converting fields to uniform units that can be averaged to form a trait value. For example, precipitation (“AA1”) was downloaded in two columns: 1) time period; 2) depth of precipitation recorded during that time period. Because the variable time periods in which data were recorded, averaging precipitation would lead to skewed results. Precipitation was changed to adapt a “precipitation per hour” model that would be more permissive to our analyses. Each trait was averaged over the time span collected (three months, one year, or three years), and the average value was designated as the phenotypic value for that strain. Only traits with values in more than 90% of strains were analyzed further.

### Association mapping

Genome-wide association (GWA) mapping was performed using 152 genome-wide *C. elegans* isotypes using the cegwas R package found at https://github.com/AndersenLab/cegwas. This package uses the EMMA algorithm for performing association mapping and correcting for population structure (Kang *et al.* 2008), which is implemented by the GWAS function in the rrBLUP package (Endelman 2011). The kinship matrix used for association mapping was generated using a whole-genome high-quality single nucleotide variant (SNV) set (Cook *et al.* 2016) and the A.mat function from the rrBLUP package. Single-nucleotide variants identified using RAD-marker sequencing (Andersen *et al.* 2012) that had at least 5% minor allele frequency in the 152 isotype set were used for performing GWA mappings. Association mappings that contained at least one SNV that had a −log(*p*-value) greater than the Bonferroni-corrected *p*-value were processed further using fine mapping, which entails a Spearman’s rank correlation test with variants from the whole-genome sequence data of moderate to severe predicted effects as determined by the SnpEff function (Cingolani *et al.* 2012).

### Temperature competition assays

We chose two strains, CX11314 and JU847, that had different alleles for the peak QTL marker (chrV: 14,822,276; JU847: T, CX11314: A) in our three-year temperature GWA mapping. JU847 has the reference allele for the peak marker and was isolated at a low temperature, whereas CX11314 has the alternative allele for the peak marker and was isolated at a higher temperature. We designed a Taqman probe (5’-[A]CCGTTTTTTTT[T/A]AATTTT-3’) to measure each of these two alleles from mixed samples of nematodes using the standard software from Applied Biosystems (https://www.thermofisher.com/order/custom-genomic-products/tools/genotyping/) and a corresponding primer set to amplify the region of interest (below).

F: 5’-AAACCCAAGATTTTTATGGTTACTTTAAGATTTGT-3’;

R: 5’-ATCTATAGTTAACTTGGATATATTGTTTGTTTTCGGT-3’

These two strains were chunked to fresh 10 cm NGMA plates seeded with OP50. 48 hours later, seven L4s from each strain were added to each of 45 6 cm NGMA plates seeded with OP50 for both 15°C and 25°C competition experiments. The 45 plates at each temperature represent nine experimental replicates each composed of five independent populations. Plates were placed at either 15°C or 25°C and grown to starvation. After one week for 25°C competition experiments or 10 days for 15°C competition experiments, nematodes were transferred to fresh NGMA plates by cutting a 0.5 cm x 0.5 cm square of agar (containing ~100 worms) and replaced at the appropriate temperature. After culture transfers two, four, and six, starved animals were washed off the plates with M9, and DNA was collected using the Qiagen DNeasy Kit. Genomic DNA from each time point was digested with the *Eco*RI enzyme and purified using the Zymo DNA Clean & Concentrator Kit. The concentration of fragmented genomic DNA was adjusted to 2 ng/µL by Qubit assay. The number of JU847 and CX11314 alleles in each replicate population was measured using Taqman analysis in a Biorad QZ200 digital droplet PCR system (File S4). Digital PCR was performed following the standard protocol provided by Biorad with the absolute quantification method. The proportion of the JU847 allele and the relative selection coefficients were calculated.

### Data Availability

Strains are available through the *Caenorhabditis elegans* Natural Diversity Resource (CeNDR, www.elegansvariation.org). File S1 contains the strains, location data, weather station data, and all traits used in mappings. File S2 contains the association mapping data. File S3 contains the interval fine mapping raw data. File S4 contains the count data from the digital droplet PCR for JU847 and CX11314 alleles.

## LITERATURE CITED

Andersen, Erik C., Justin P. Gerke, Joshua A. Shapiro, Jonathan R. Crissman, Rajarshi Ghosh, Joshua S. Bloom, Marie-Anne Félix, and Leonid Kruglyak. 2012. “Chromosome-Scale Selective Sweeps Shape Caenorhabditis Elegans Genomic Diversity.” Nature Genetics 44 (3): 285–90.

Bozinovic, Francisco, Nadia R. Medina, José M. Alruiz, Grisel Cavieres, and Pablo Sabat. 2016. “Thermal Tolerance and Survival Responses to Scenarios of Experimental Climatic Change: Changing Thermal Variability Reduces the Heat and Cold Tolerance in a Fly.” Journal of Comparative Physiology. B, Biochemical, Systemic, and Environmental Physiology 186 (5): 581–87.

Cingolani, Pablo, Adrian Platts, Le Lily Wang, Melissa Coon, Tung Nguyen, Luan Wang, Susan J. Land, Xiangyi Lu, and Douglas M. Ruden. 2012. “A Program for Annotating and Predicting the Effects of Single Nucleotide Polymorphisms, SnpEff: SNPs in the Genome of Drosophila Melanogaster Strain w1118; Iso-2; Iso-3.” Fly 6 (2): 80–92.

Cook, Daniel E., Stefan Zdraljevic, Robyn E. Tanny, Beomseok Seo, David D. Riccardi, Luke M. Noble, Matthew V. Rockman, et al. 2016. “The Genetic Basis of Natural Variation in Caenorhabditis Elegans Telomere Length.” Genetics, July. doi:10.1534/genetics.116.191148.

Endelman, J. B. 2011. “Ridge Regression and Other Kernels for Genomic Selection with R Package rrBLUP.” The Plant Genome. dl.sciencesocieties.org. https://dl.sciencesocieties.org/publications/tpg/abstracts/4/3/250.

Félix, Marie-Anne, and Christian Braendle. 2010. “The Natural History of Caenorhabditis Elegans.” Current Biology: CB 20 (22): R965–69.

Frézal, Lise, and Marie-Anne Félix. 2015. “C. Elegans Outside the Petri Dish.” eLife 4 (March). doi:10.7554/eLife.05849.

Gerken, Alison R., Olivia C. Eller, Daniel A. Hahn, and Theodore J. Morgan. 2015. “Constraints, Independence, and Evolution of Thermal Plasticity: Probing Genetic Architecture of Long- and Short-Term Thermal Acclimation.” Proceedings of the National Academy of Sciences of the United States of America 112 (14): 4399–4404.

Hangartner, Sandra B., Ary A. Hoffmann, Ailie Smith, and Philippa C. Griffin. 2015. “A Collection of Australian Drosophila Datasets on Climate Adaptation and Species Distributions.” Scientific Data 2 (November): 150067.

Hijmans, R. J., E. Williams, and C. Vennes. 2012. “Geosphere: Spherical Trigonometry. R Package Version 1.2–28.” CRAN. R-Project. Org/package= Geosphere.

Hirschhorn, Joel N., and Mark J. Daly. 2005. “Genome-Wide Association Studies for Common Diseases and Complex Traits.” Nature Reviews. Genetics 6 (2): 95–108.

Hodgkin, J., and T. Doniach. 1997. “Natural Variation and Copulatory Plug Formation in Caenorhabditis Elegans.” Genetics 146 (1): 149–64.

Hutchinson, G. E. 1957. “Cold Spring Harbor Symposium on Quantitative Biology.” Concluding Remarks.

Iannone, Richard. 2015. “stationaRy: Get Hourly Meteorological Data from Global Stations. R Package Version 0.4.1.” CRAN. R-Project. Org/package= stationaRy, October. Comprehensive R Archive Network (CRAN).

“Integrated Surface Global Hourly Data - NOAA Data Catalog.” 2016. Accessed September 2. https://data.noaa.gov/dataset/integrated-surface-global-hourly-data.

Kang, Hyun Min, Noah A. Zaitlen, Claire M. Wade, Andrew Kirby, David Heckerman, Mark J. Daly, and Eleazar Eskin. 2008. “Efficient Control of Population Structure in Model Organism Association Mapping.” Genetics 178 (3): 1709–23.

Li, Heng, and Richard Durbin. 2009. “Fast and Accurate Short Read Alignment with Burrows–Wheeler Transform.” Bioinformatics 25 (14): 1754–60.

Li, Heng, Bob Handsaker, Alec Wysoker, Tim Fennell, Jue Ruan, Nils Homer, Gabor Marth, Goncalo Abecasis, Richard Durbin, and 1000 Genome Project Data Processing Subgroup. 2009. “The Sequence Alignment/Map Format and SAMtools.” Bioinformatics 25 (16): 2078–79.

Machado, Heather E., Alan O. Bergland, Katherine R. O’ Brien, Emily L. Behrman, Paul S. Schmidt, and Dmitri A. Petrov. 2016. “Comparative Population Genomics of Latitudinal Variation in Drosophila Simulans and Drosophila Melanogaster.” Molecular Ecology 25 (3): 723–40.

Saltz, Julia B., and Sergey V. Nuzhdin. 2014. “Genetic Variation in Niche Construction: Implications for Development and Evolutionary Genetics.” Trends in Ecology & Evolution 29 (1): 8–14.

Sterken, Mark G., L. Basten Snoek, Jan E. Kammenga, and Erik C. Andersen. 2015. “The Laboratory Domestication of Caenorhabditis Elegans.” Trends in Genetics: TIG 31 (5): 224–31.

Taylor, Michelle L., Alison Skeats, Alastair J. Wilson, Tom A. R. Price, and Nina Wedell. 2015. “Opposite Environmental and Genetic Influences on Body Size in North American Drosophila Pseudoobscura.” BMC Evolutionary Biology 15 (March): 51.

Tyukmaeva, Venera I., Paris Veltsos, Jon Slate, Emma Gregson, Hannele Kauranen, Maaria Kankare, Michael G. Ritchie, Roger K. Butlin, and Anneli Hoikkala. 2015. “Localization of Quantitative Trait Loci for Diapause and Other Photoperiodically Regulated Life History Traits Important in Adaptation to Seasonally Varying Environments.” Molecular Ecology 24 (11): 2809–19.

